# Figbird: A probabilistic method for filling gaps in genome assemblies

**DOI:** 10.1101/2021.11.24.469861

**Authors:** Sumit Tarafder, Mazharul Islam, Swakkhar Shatabda, Atif Rahman

## Abstract

**Motivation:** Advances in sequencing technologies have led to sequencing of genomes of a multitude of organisms. However, draft genomes of many of these organisms contain a large number of gaps due to repeats in genomes, low sequencing coverage and limitations in sequencing technologies. Although there exist several tools for filling gaps, many of these do not utilize all information relevant to gap filling.

**Results:** Here, we present a probabilistic method for filling gaps in draft genome assemblies using second generation reads based on a generative model for sequencing that takes into account information on insert sizes and sequencing errors. Our method is based on the *expectation-maximization* (EM) algorithm unlike the graph based methods adopted in the literature. Experiments on real biological datasets show that this novel approach can fill up large portions of gaps with small number of errors and misassemblies compared to other state of the art gap filling tools.

**Availability and Implementation:** The method is implemented using C++ in a software named “Filling Gaps by Iterative Read Distribution (Figbird)”, which is available at: https://github.com/SumitTarafder/Figbird.

**Supplementary information:** Supplementary data are available at *Bioinformatics* online.

## 1 Introduction

Genome sequencing of an organism i.e. determining the sequence of nucleotides that make up the entire genome is often a prerequisite for performing experiments to study that organism and fundamental to understanding how different organisms relate to each other. With the advancement of sequencing technologies in the past decade and a half, the availability of sequencing data has increased drastically and genomes of many organisms have been sequenced. However, genomes can be billions of base pairs long and there is still no technology that can sequence entire genomes at one go. So instead of sequencing the complete genome, different sequencing technologies generate millions of smaller fragments called reads. The whole genome is then reconstructed from these read sequences through a process known as genome assembly.

The high throughput, low cost and error rates of second generation technologies such as Illumina [1], have led to its use in a large number of sequencing projects and many assembly tools such as ABySS [2], Velvet [3], Allpaths-LG [4] etc. have been developed to sequence genomes using this technology. However, due to the short lengths of second generation reads, assembly is performed in multiple steps. The first step of the assembly pipeline constitutes of stitching the read sequences into contiguous sequences called contigs and then in the next step paired-end or mate pair reads are used to orient and organize the contigs into scaffolds. Despite the development in methodologies for genome assembly, the draft assemblies constructed with these tools still contain thousands of intervening gaps within the assembled scaffolds due to repetitive regions in genomes and regions with low sequencing coverage. Filling these gaps with minimal introduction of error is a crucial step in genome assembly pipeline as an error free complete genome can lead to better downstream analysis such as genotyping variants without error, complete annotation of genes [5], identification of effector and co-regulated genes [6], and accurate statistical analysis [7].

To address this issue, a number of tools have been developed for gap filling using short reads. Many of the genome assemblers such as ABySS and Allpaths-LG include gap-filling modules in their pipeline. In addition, stand-alone tools such as GapCloser in SOAPdenovo[8]package, GapFiller [9], Gap2Seq [10], Sealer[11], etc. also exist for this purpose. GapCloser constructs a De Bruijn graph on the set of available reads to perform the local assembly. Although it works well for smaller genomes, it is highly memory inefficient [11] for larger genomes and only considers read pairs with insert size less than 2000 base pair. GapFiller on the other hand uses read pairs with one end aligned to the scaffold and the other end partially aligned to gap regions. The reads are then assembled using a k-mer based method to fill the gaps. A major limitation of GapFiller is that it does not use a lot of sequence information due to the use of partial reads only. A recent computational approach for gap filling has been introduced in [10] where the problem is formulated as an exact path length (EPL) problem, implemented in pseudo-polynomial time with some optimizations and packaged in a tool called Gap2Seq. However, their approach does not scale to large genomes and is unable to fill large gaps due to the expensive computational approach. Lastly, a resource efficient gap-filling software named Sealer [11] has been designed to close gaps in scaffolds by navigating De Bruijn graphs represented by space-efficient bloom filter data structures and is claimed to be scalable to large giga base pair sized genomes. It uses an assembly utility within the ABySS package, called Konnector [12] as its engine to close intra-scaffold gaps. The Konnector utility takes the flanking sequence pairs along with a set of reads with a high level of coverage redundancy as inputs and runs with a range of k-mer lengths to connect the flanking gap sequences. Sealer ignores size discrepancies between gaps and newly introduced sequences, since gap sizes are often estimated from fragment library distributions and assemblers do not generally provide confidence intervals for every region of Ns, but does not consider the insert size of paired reads during gap filling.

Almost all of the methods for gap filling use a graph based formulation of the problem, most commonly De Bruijn graph, and then try to find an Eulerian path through the graph that corresponds to the gap sequence. But there can be multiple such paths present in the graph due to repetitive regions or sequencing errors [3] and only one of those paths corresponds to the true genomic sequence of the gap. Finding such a correct path can get complicated either due to the presence of repeats or because of the memory constraint due to the nature of the graph built from a large set of k-mers. Moreover, once the reads to be used for filling a gap are identified, most of the tools ignore distance information from the other end of the pair i.e. insert size, that may help disambiguate among multiple sequences and solve repeat related issues. So, searching for the true gap sequence that can solve the above stated problems is still an area to be explored in genome assembly.

Recently, sequencing technologies have gone through a further revolution and “third generation” single molecule technologies, such as Pacific Bio-sciences and Oxford Nanopore have been developed which can generate read sequences of lengths tens to hundreds of kilo-base pairs (Kbp) and beyond. Among the long read and/or contig based gap closing approaches existing in literature, GMcloser [13], PBJelly [14] and gapFinisher [15] are worth mentioning which use long reads or alternatively assembled contig set from short read libraries to fill gaps using sequence alignment and consensus building procedure. Although long reads lead to substantially better genome assemblies [16], its use in the gap filling process is still limited due to the high error rate in these technologies. Both PacBio and Nanopore technologies have far higher error rate (88 – 94% accuracy for Nanopore and 85 – 87% for PacBio) as opposed to 99.9% accuracy for second generation Illumina HiSeq 4000 [17]. To mitigate the effect of error-prone data, high coverage is required to ensure low error rate in the generated assemblies. As gap filling is the last stage of genome assembly pipeline, any error introduced in this stage will carry over subsequent genomic analysis and may lead to incorrect results. To this end, we have chosen comparatively accurate second generation read sequences for gap filling purposes.

In this work, we formulate gap filling as a parameter estimation problem and develop a probabilistic method for filing gaps in scaffolds using second generation reads. The method is based on a generative model for sequencing proposed in CGAL [18] and subsequently used to develop a scaffolding tool SWALO [19]. The model incorporates information such as distribution of insert size of read pairs, sequencing errors, etc. and can be used to compute likelihood of an assembly with respect to a set of read pairs. We use this model to estimate the length of the gap and to find a sequence for each gap that maximizes probability of the reads mapping to that gap region. However, in this case the insert sizes of the read pairs that have only one end mapped is unknown. So, we use an iterative approach based on the *expectation-maximization* (EM) algorithm [20], which has been used to solve many problems in computational biology including motif finding [21] and transcript abundance estimation from from RNA-Seq [22]. Our method is implemented in a tool called Figbird and an extensive comparison with other standalone gap fillers on data sets from the GAGE [23] project is performed. Overall, our probabilistic method performs well consistently on six different metrics over variety of real draft assemblies, and is able to reduce amount of gaps substantially while keeping missassemblies and errors low. Although the method is computationally intensive, it is linear in the number of reads as well as the number and the lengths of gaps, and is thus scalable and applicable to large genomes.

## 2 Methods

In this section, we describe the methods behind Figbird. First, we present an overview of the method and subsequently discuss each step in detail.

### 2.1 Overview of Figbird

A high level overview of our gap-filling method is illustrated in the block diagram in Figure 1. The method consists of the following two major phases.

**Fig. 1:**
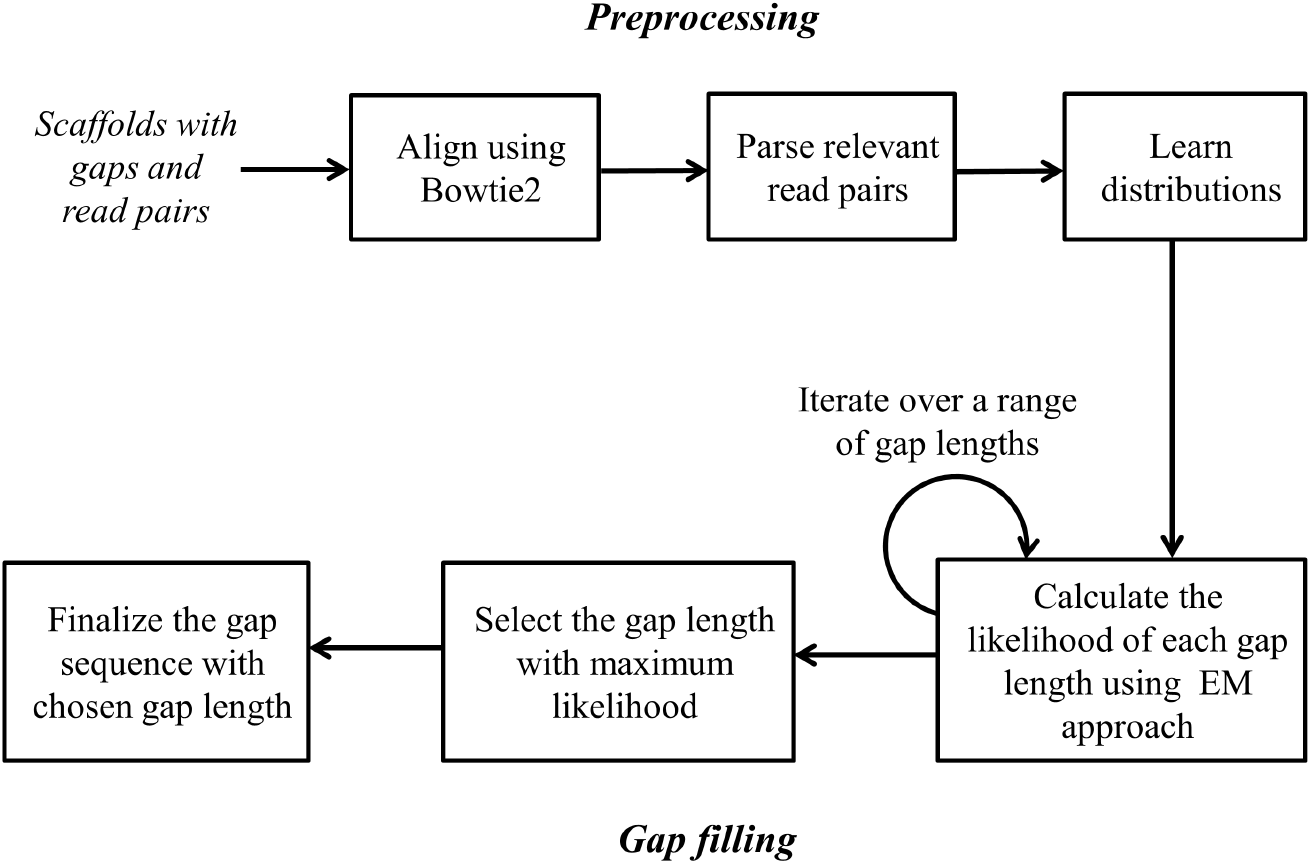
Overview of Figbird. Figbird takes as input scaffolds containing gaps and sets of read pairs (paired end or mate pairs) and aligns the reap pairs to the scaffolds using Bowtie2. The output is then parsed to separate fully mapped as well as to assign one end unmapped and one end partially mapped read pairs to specific gaps. Insert size and error distributions are then learnt using uniquely and fully mapped read pairs. In the gap filling phase one end unmapped and partially mapped reads are used to fill gaps using the expectation-maximization (EM) algorithm for a range of gap lengths. Finally, reads with probabilities below a threshold are filtered and consensus sequence is formed for the gap length with maximum likelihood

#### Pre-processing

In this phase, relevant read pairs are identified and distributions are learned. At first paired end and mate pair reads (we will refer to both these types as read pairs) are aligned to the scaffold set using Bowtie2 [24]. Then the output of Bowtie2 is parsed to collect relevant read pairs necessary for our method. The details of the read pairs as well as the read parsing criteria is described in Subsection 2.2. Next, the insert size distribution and parameters for our error model are learned using uniquely and fully mapped read pairs.

#### Gap filling

In the gap filling phase, read pairs with one end unmapped or partially mapped are locally assembled using a maximum likelihood approach calculated using a model described in Subsection 2.4. As we do not know exactly where the unmapped end of the read pair should be placed within the gap, we use the *expectation-maximization(EM)* algorithm [20] to iteratively find the placement of the read using the learned insert size distribution and the current estimate of the nucleotides in the gap sequence, and re-estimate the probabilities of the nucleotides using the current placements. The process is iterated over a range of gap estimates with different lengths and the one with the maximum likelihood value is chosen to fill the gap region. Once the gap length and the corresponding sequence is estimated, the distribution of probability of fully mapped reads learnt from the previous step are used to decide whether the reads should be considered to fill that particular gap based on a cut-off value and reads with probability below the cut-off are discarded. Finally, a consensus is calculated based on the probabilistic placements of the chosen reads in gaps which is regarded as the final predicted gap sequence.

### 2.2 Aligning and parsing read pairs

In this phase, read pairs are aligned to the draft scaffolds using Bowtie2 and the resulting output file in SAM format [25] is parsed to separate read pairs, and to assign relevant reads to specific gaps. The different types of reads considered are shown on Figure 2 and described below:

**Fig. 2:**
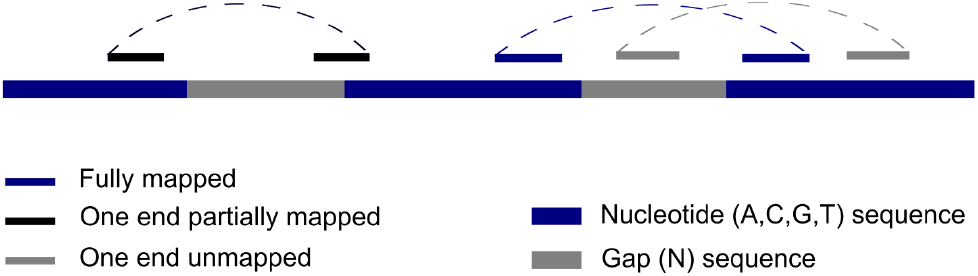
Different types of read pairs used for gap filling.

#### (i) Fully aligned read pairs

Fully aligned read pairs are the ones whose both ends map to the draft genome concordantly i.e. in proper orientation and within the specified insert size. These type of reads are used to learn insert size distributions and sequencing error rates which will be used to calculate the likelihood of a gap estimate.

#### (ii) One end unmapped pairs

These are the read pairs which have one end mapped to the draft scaffold but the other end remains unmapped. To find which gap the unmapped end is possibly associated with, the mapped end of the read pair is checked to determine whether it falls within 1.15 × *M* base pairs on either side of the gap region and these pairs are then separately saved per gap region. Here, *M* denotes the insert size mean of the corresponding library given input to Figbird and the threshold is set based on the observation that most of the insert sizes fall within one hundred and fifteen percent of the input mean.

If the condition is satisfied for more than one gap due to the congested and fragmented nature of the gaps, then it is ambiguous as to which gap the unmapped read actually belongs to. In that case, we chose the gap for which the distance is closest in terms of insert size mean of the library.

#### (iii) One end partially mapped pairs

By running Bowtie2 in the mode that supports local alignment, we also collect the read pairs whose one end is partially mapped to one of the ends of a gap region. These type of reads should lead to an error free construction of gap sequences as the placement of the reads are known in this case.

Due to the potential presence of huge number of gaps as well as the large lengths of those gaps, a lot of read pairs often have none of their ends mapped to the draft genome. Since our method will be run several times on different read sets, there is a chance that the reads that are unmapped in initial iterations will be mapped in subsequent iterations. So in addition to using the three types of reads stated above, we may also be able to utilize both end unmapped reads as well for gap filling.

### 2.3 Learning distributions

In order to fill gaps using Figbird, we need to learn insert size distribution and sequencing error characteristics which are required to calculate the probability that a set of read pairs is generated from a certain region of the genome using the generative model presented in CGAL [18]. Since sequencing characteristics differ among different libraries, we have chosen to learn distributions from individual libraries used in the experiments. To do that, we map the read pairs to the scaffolds using Bowtie2 and learn empirical distributions using fully mapped read pairs that map uniquely. As the number of uniquely mapped reads to learn the insert size distribution from may not be very high for some datasets, it is smoothed using a window size of 12 as well as truncated as proposed in [19]. We calculate the insert size distribution for each dataset as shown in examples in Supplementary Figure S1. We also compute the mean *μ* and left-sided and right-sided standard deviations, *σ_l_* and *σ_r_* respectively and truncate the distribution at *μ* – 2.5*σ_r_* and *μ* + 2.5*σ_r_* which we use as our minimum and maximum threshold for the insert size range of an unmapped read. The error model used in our computation, described in detail in [18], includes substitutions or mismatches, insertions-deletions and takes into account differences in error rates across positions in reads.

### 2.4 Gap filling using the EM algorithm

In the next phase, we fill the gaps using read pairs with one end mapped and the other end unmapped or partially mapped with a likelihood based approach. Given a gap sequence 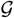 of length *g* and a set of unmapped reads 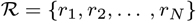, the log likelihood of 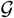 is given by

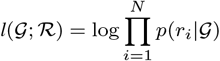

where 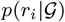 is the probability that *r_i_* is generated from 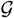. We are interested in the gap sequence that maximizes this likelihood and to calculate this, we use the generative model described in CGAL. However, since one end of the read pair is mapped to a fixed position, we modify the model and define probability of a read as follows:

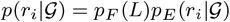

where *L* is the insert size of the read pair, and *p_F_* and *p_E_* are insert size and error distributions respectively.

In this paper, we formulate gap filling as a parameter estimation problem. Given a gap of length *g*, we introduce the parameter to be estimated as

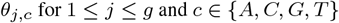

where *θ_j,c_* denotes the probability of nucleotide *c* at index *j* of gap sequence. The gap filling problem then converts into a parameter estimation problem where the goal is to find the estimates of *θ_j,c_*s that maximize the likelihood 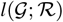.

However, to estimate these parameters, we need to know the insert sizes exactly. Although the insert sizes are known for one end partially mapped reads, we do not know these for one end unmapped reads as sequencing experiments do not provide the exact distance between the two ends. It is worth noting that although we don’t exactly know the insert size values, the read pairs follow an approximately normal distribution which can be learnt as discussed earlier and this observation can reduce the possible positions of the unmapped end within a range of minimum and maximum value of insert size for that read pair as indicated in Figure 3.

**Fig. 3:**
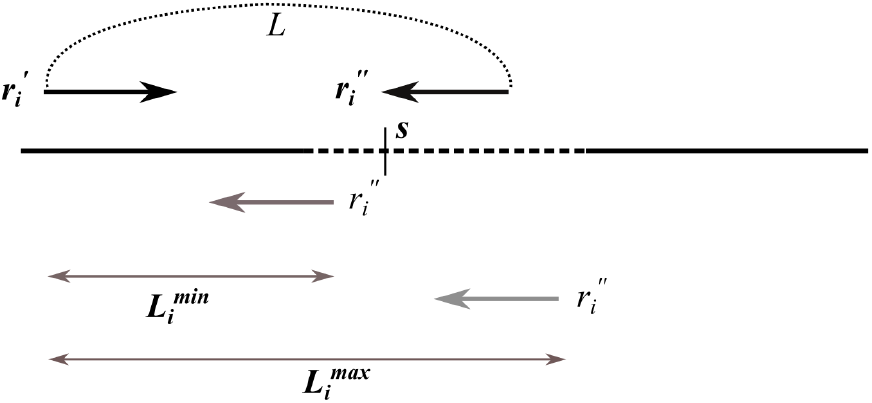
Possible placement of the unmapped end 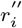 of read *r_i_* in different gap positions.

Now, if the insert size of a read pair was exactly known, we could have placed the read at the correct position within the gap and use the sequence to adjust the probabilities of the nucleotides occurring at those gap positions as shown in Figure 4A. On the other hand, if the gap sequence was known to us beforehand, then we could have aligned the read to that known sequence and obtain the likely placement of the unmapped end as shown in Figure 4B.

**Fig. 4:**
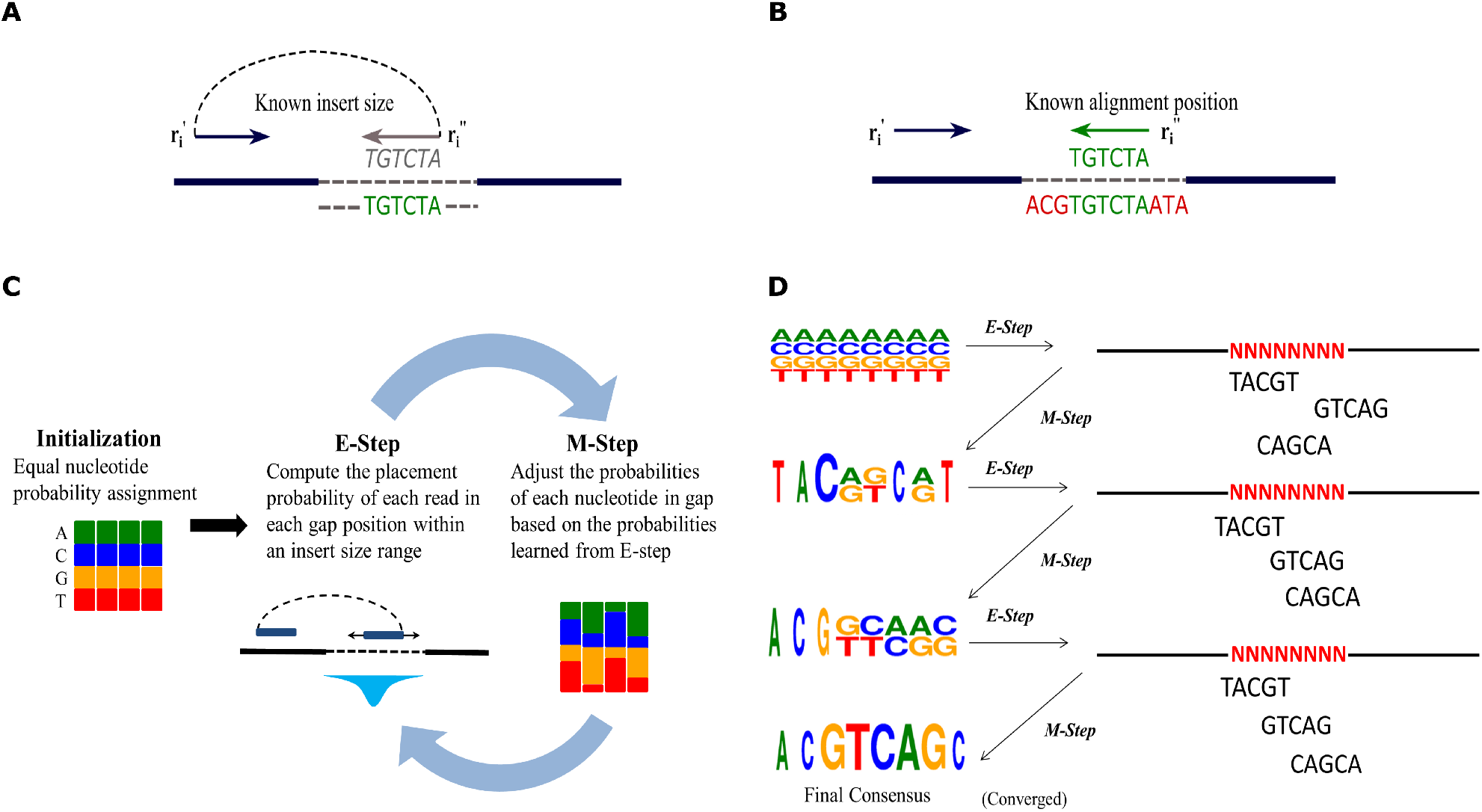
Formulation of gap filling using the EM algorithm. (A) If the insert sizes are known exactly, the reads can be placed within the gap in the correct positions and the gap sequence can be inferred. (B) If the sequence of nucleotides in the gap is known, reads can be aligned to the sequence, and their placement and insert sizes can be estimated. (C) Figbird solves gap filling using the EM algorithm. It starts by initializing each nucleotide with equal probability. In the E-step, current probabilities of nucleotides and insert size distribution is used to calculate placement probabilities of each read within the gap. Then in the M-step, the placement probabilities and read sequences are used to update the probabilities of nucleotides. These two steps are iterated until convergence. (D) A simplified simulation of the EM algorithm for gap filling.

But neither the exact insert sizes nor the sequence of the gap region are known beforehand and to solve one of these problems we need the solution of the other. To solve this set of interlocking problems, we will use the *expectation maximization* (EM) algorithm. EM algorithm can be applied to solve those class of problems which have some hidden observations as well as unknown model parameters. It proceeds by picking an initial set of model parameters to estimate the hidden observations with the assumption that the data comes from a specific model. This is called the *E-step.* Then, using the newly estimated values of hidden observations, the parameters or initial hypothesis gets updated. This step is called the *M-step.* These two steps are iterated until the resulting values converge to a fixed point or the allocated time ends. In our method, the hidden observations are insert sizes of the read pairs and the parameters are the probabilities of each nucleotide occurring at each gap position.

#### EM formulation

A schematic diagram of our *expectation maximization* approach for gap filling is shown in Figure 4C. We start with an initial hypothesis that each of the four bases {*A, C, G, T*} are equally probable at each gap position *j* and initialize the estimation parameter *θ_j,c_* with 0.25 for all gap positions and for all four possibilities of nucleotide *c*. Then the E step and M step will be applied iteratively as follows:

##### E-step

In the E-step, we place the unmapped end of the read *r_i_* = *c*_1_ *c*_2_ … *c_l_* in all possible gap positions based on the set of allowable insert sizes 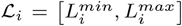, where 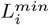 and 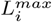 are the minimum and maximum threshold value of insert size for read *r_i_* respectively and calculate the probability that the read is generated from that position using the current estimated probabilities *θ*. This posterior probability of read *r_i_* having a particular insert size 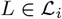, i.e. that it starts at *s,* is given by:

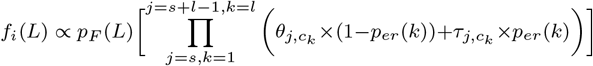

where *l* is the length of the read and *s* is the start position of the read within the gap corresponding to the insert size *L, p_er_*(*k*) is the error probability at read position *k,* and *τ_j,c_k__* is the probability of getting nucleotide *c* at gap index *j* and read position *k* due to a sequencing error which can be calculated using the following equation:

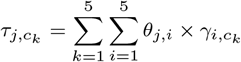

where, *γ* is a 5 × 5 square matrix containing the probability of substitution of each of the four nucleotide characters and character ‘N’ into others.

##### M-step

In this step, we accumulate the probabilities calculated for all the reads in E-step in an intermediate matrix *α* with the same dimensions as *θ*. If s is the starting position of read 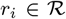 in the gap with respect to an insert size 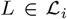, *α* will be updated according to the following equation:

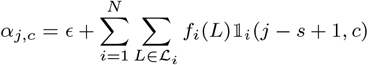

where *ϵ* is a small value added to ensure the probability of characters do not become zero and 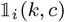 is a variable indicating whether the *k*-th character *of r_i_* equals *c* i.e.

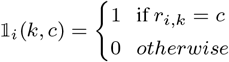

Finally, the update of our estimation parameter *θ* using the intermediate values in *α* will be done as following:

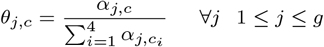

Here, *c_i_* denotes each of the four possible nucleotides in position *j*. Based on this updated hypothesis, we will continue our E and M steps until there is a convergence i.e. the placement positions of reads don’t change anymore and thus the hypothesis reaches a fixed set of values.

A simplified simulation of our EM algorithm is presented in Figure 4D. Initially, the probabilities of each nucleotide is equal across all positions as indicated in the top left of the figure. Based on this set of parameters, we calculate the probabilities of each read aligning at each gap position in the E-step and accumulate those probabilities at M-step to determine an intermediate consensus with updated nucleotide probabilities. The relative height and frequency of nucleotides at different positions in the figure denote the corresponding information content and relative probabilities at those positions. These two steps then iterate two more times and finally at the end of third iteration we manage to obtain the true placements of the unmapped reads in gap based on the continuously updated set of parameters and the steps converge to a fixed value. The final consensus is constructed based on the final placement of reads using a majority voting approach and the gap is filled with the predicted final consensus sequence.

### 2.5 Selecting the gap length

Once the EM converges for a particular gap length *g*, we compute the likelihood of the estimated parameters as follows:

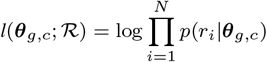

where *p*(*r_i_*|*θ_g,c_*) is the maximum over all placement probabilities of *r_i_* i.e.

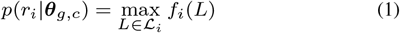

However, since the gap length is often not known exactly, we iterate over a range of gap lengths and select the gap length that maximizes the above likelihood i.e. 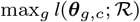. For each gap with length *g* in the scaffold file, we compute the likelihood for the range 0.5 × *g* to 2.5 × *g* and the gap estimate with the maximum likelihood will be selected as the final gap estimate. We observe that for most of the gaps, the actual gap length is within the specified range. However, for long gaps this becomes computationally expensive. So, we use a heuristic for deciding the range which is described in Supplementary Note 3.2. It is to be noted that gap length can be negative if the sequences preceding and succeeding the gap region overlaps.

### 2.6 Finalizing the gap sequence

At this stage of our pipeline, we will finalize our gap sequence based on an error model and learnt cut-off value. The significance of this step is that every unmapped read parsed during the preprocessing phase does not truly belong to that gap region due to the fact that one or both end of a read pair might be very error prone and the aligner sometimes fail to map these reads to the scaffolds and thus they remain unmapped. So it is essential not to consider such reads in gap filling process. To prune these unnecessary reads out, we perform the following steps. Firstly, we generate an intermediate consensus sequence 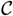 based on the output of final M-step of our method for the gap length corresponding the highest likelihood. Then each unmapped read *r_i_* will be slid across the allowable consensus positions based on the insert size, and the error probability of *r_i_* placed on consensus position *s* will be calculated using the following equation:

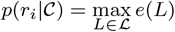

If the consensus character at position *j* i.e 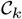 matches with *r_ij_* which is the *j^th^* character of read *r_i_*, then

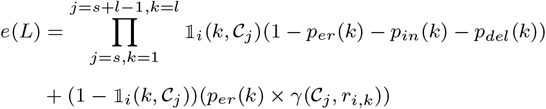

where *p_er_, p_in_, p_del_* are error position, insertion and deletion distributions respectively across all read position calculated using the parsed CIGAR information from SAM alignment output. If the error probability 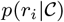 if *r_i_* is less than a cut off error probability, only then we will consider the read for gap filling, otherwise it will be discarded from consensus consideration. The cut off probability value is pre-calculated using the distribution of probabilities of uniquely and fully mapped reads. It is the value above which the probabilities of 80% of such reads lie.

Finally, we place each read that are above the cut-off threshold value at their most likely position according to Equation 1 and construct a final consensus sequence based on a maximum voting approach. This will be our final predicted sequence 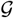 for that particular gap.

### 2.7 Implementation

The method is implemented using C++ and is available for download freely at: https://github.com/SumitTarafder/Figbird. In order to reduce the runtime and ensure accuracy, we apply the following heuristics in the implementation:

- **Removal of duplicate reads:** We have removed duplicate reads while parsing reads with one end unmapped which helps significantly in the calculation of likelihood due to the presence of erroneously parsed reads as discussed in Subsection 2.6. We have observed that such reads tend to pile up in certain parts of gaps due to the sequence similarity and including them negatively affects both the accuracy and running time of the method.
- **Iterative implementation:** To make the most out of the various types of reads available, we run Figbird with one end partially mapped and one end unmapped reads of different libraries in an iterative manner. A detailed description of the iterations are given in Supplementary Note 3.1.
- **Reduction of read library:** To make the method scalable, we have removed read pairs with both ends mapped from our dataset after the first iteration as learning the distributions and error parameters once suffices for the rest of the iterations.
- **Other heuristics:** To improve accuracy in the predicted consensus sequence, a number of heuristics are applied at different stages of gap filling which are described in Supplementary Notes 3.2:

## 3 Results

### 3.1 Experimental data and setup

To assess the performance of Figbird and to compare it with existing gap filling tools, we use the GAGE dataset [23]. It is a standard dataset that was generated to critically evaluate the genome assemblers and is also used to assess gap filling tools [10]. In this experiment, we use the data for two bacterial species *Staphylococcus aureus, Rhodobacter sphaeroides* as well as *Homo sapiens* Chromosome 14, as shown in Table 1, for which reference genomes are available. We collect a wide array genome assemblies for the three datasets generated using various tools as part of the GAGE project and performed gap filling using Figbird on each of them, and evaluate the results by comparing the filled sequences with the reference sequences.

**Table 1.** Genomes used in evaluation

| Organism | Genome size | No. of assemblies | No. of gaps | Gap length range |
| --- | --- | --- | --- | --- |
| *Staphylococcus aureus* | 2 - 3 Mbp | 8 | 35 - 654 | 1 - 3462 |
| *Rhodobacter sphaeroides* | 3 - 5 Mbp | 9 | 38 - 938 | 1 - 4808 |
| Human Chromosome 14 | 80 - 150 Mbp | 9 | 1061 - 51567 | 1 - 34745 |

As part of the GAGE datasets, second generation sequencing reads from two libraries are available for each of these three datasets. For our experiment, we use both the fragment (paired-end) as well as the short jump (mate pair) libraries. The details about the short reads libraries used in this experiment are listed in Table 2. For more details about the different assemblies and read sets, readers are encouraged to check the official GAGE website. For sequences from the short jump library, we use Quake [26] corrected versions of the reads for better accuracy due to it’s conservative nature of error correction mechanism [27]. We run Bowtie2 version 2.2.3 to align these read pairs to the gapped scaffolds for our experiment and then fill the gaps using Figbird.

**Table 2.** Read libraries used in evaluation

| Organism | Sequence library | Mean insert size | Read length | No. of read pairs |
| --- | --- | --- | --- | --- |
| *Staphylococcus aureus* | Frag | 180 | 101 | 1,294,104 |
|  | Shortjump | 3500 | 30 - 37 | 1,614,660 |
| *Rhodobacter sphaeroides* | Frag | 180 | 101 | 2,050,868 |
|  | Shortjump | 3500 | 76 - 101 | 1,526,850 |
| Human Chromosome 14 | Frag | 155 | 101 | 36,504,800 |
|  | Shortjump | 2283 - 2803 | 70 - 101 | 14,054,994 |

We compare the performance of our tool Figbird with the four state-of-the-art tools available for filling gaps using short reads which are, SOAPdenovo’s stand-alone tool GapCloser v1.12-r6 [8], GapFiller v1.10

[9], Gap2Seq v1.0 [10] and Sealer [11]. For GapFiller, both BWA [28] and Bowtie [29] aligners are used.

All experiments are run using 24 cores on a machine with Intel(R) Xeon(R) CPU E5-2697 v2 @ 2.70GHz processors. We used the time.h library in C++ to measure the time taken for each of these tools and used the Python script Memusg to benchmark peak memory usage.

### 3.2 Evaluation criteria

As gap-filling is one of the last steps in the genome assembly pipeline, a wrongly introduced sequence may affect subsequent analysis especially if the gap region falls in coding regions of the genome. Therefore, we focus on errors introduced by gap filling tools in addition to the amounts of gaps filled by them. We use QUAST v2.3 [30] to compare the outputs of the tools with the reference, and then use a Python script provided by [10] for analysis of the results and for classification into “misassemblies” and other errors. QUAST uses NUCmer [31] to find alignments between the gap-filled assembly and the reference sequence.

To assess the quality of the filled sequence and the robustness of the methods, we list six different metrics described below:

- **Misassemblies**: The number of misassembled sequences in a scaffold that are larger than *M* bp. Here, *M* is chosen to be 4000 as in [10], which is the upper bound on the insert size of the mate pair libraries used in this experiment.
- **Erroneous length**: It is the sum of the lengths of all mismatches,, indels and local misassemblies i.e. the length of the misplaced sequence is ≤ *M*.
- **Unaligned length**: The total length of an assembly that is unaligned with the reference.
- **NGA50**: NG50 is the size of the longest scaffold such that at least half of the reference genome is contained by scaffolds longer than it. NGA50 is the NG50 after scaffolds have been broken at every position where a local misassembly or misassembly has been found.
- **Number of gaps**: The total number of gaps i.e a contiguous sequence of ‘N’ remaining where gap length is ≥ 1
- **Total gap length**: The sum of remaining amount of unknown nucleotide positions denoted by ‘N’ in the filled assembly.

### 3.3 Comparison with gap filling tools

In this section, we present the performance comparison of Figbird with four other state-of-the-art gap filling tools that use short reads as discussed in Section 3.1. The detailed evaluation results from QUAST are provided in Supplementary Tables S1-S3. Each table shows the percentage increase or decrease in the six evaluation metrics achieved by the five gap filling tools along with the original values before gap filling. For each assembly, these relative percentages are determined using the differences in evaluated values between original assembly and gap filled assembly using QUAST. The results over all the assemblies are presented in the bottom of the tables named ‘Total’, which is obtained by determining the average for a particular metric over all the assemblies for each gap filling tool. The overall percent reduction in gap length as well as percent changes in misassembly and errors for all three datasets are also shown in Figure 5. We focus on these three metrics as unaligned length and NGA50 are related to these while filling a gap partially lead to an increase in number of gaps which we believe is misleading. From the overall results, we observe that, Figbird is able to close considerably high portion of gap regions yet managing to keep the erroneous length and misassembly low compared to the other state of the art tools.

**Fig. 5:**
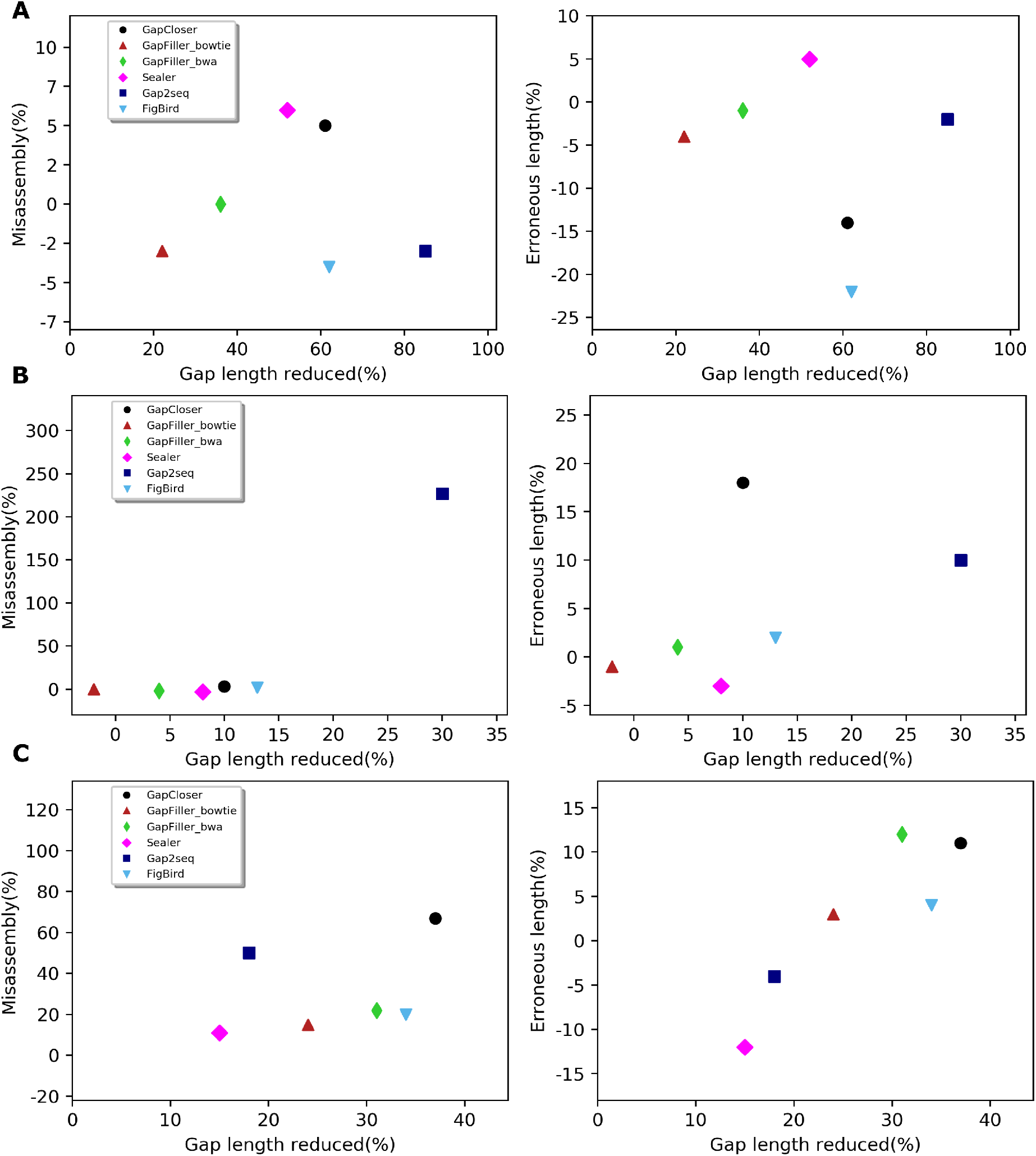
Scatter plots showing the performance of Figbird compared to other gap filling tools on (A) *S.aureus,* (B) *R. sphaeroides* and (C) Human Chromosome 14 datasets in terms of average percentage of misassemblies and erroneous length against the average percentage of nucleotides filled by each tool. The values are computed by determining the percent change in misassembly and erroneous length, and percent reduction in gap length obtained by each tool for all the de novo draft genome assemblies from GAGE, and then averaging over the values for all those assemblies.

For the *Staphylococcus Aureus* dataset, we observe from the scatter plots in Figure 5A that none of the other tools have managed to fill more nucleotides than Figbird while achieving better accuracy in terms of error or misassembly i.e. there are no points in the scatter plots that are to the right and below the points representing Figbird. We can also see from Supplementary Table S1 that Figbird has reduced the total misassembly rate by 4%, which is the best among all other tools, and reduced erroneous length by 22% thus outperforming the next best tool GapCloser in this respect by 8%. The only tool which is able fill a higher portion of gaps than Figbird is Gap2Seq which is at the expense of higher rate of misassembly and substantially higher amount of erroneous sequences.

In case of our second bacterial dataset *Rhodobacter Sphaeroides*, a similar trend can be observed from Figure 5B like the previous dataset. None of the other tools have outperformed Figbird which has managed to close second highest amount of ‘N’s, only behind Gap2Seq. However, Gap2Seq increases misassembly and erroneous length by 227% and 10% respectively which are substantially higher than the 2% misassembly and 2% erroneous length increase by Figbird, as shown in Supplementary Table S2. It is worth highlighting that the sequencing error rate for *R. sphaeroides* dataset is higher compared to that of *S. aureus* dataset and the sequencing reads being more error-prone, mapping tools finds it difficult to align reads properly to the scaffolds [32], and also leads to errors during gap filling. This shows that Gap2Seq lacks robustness to sequencing errors and may introduce large amount of errors whereas Figbird is still performs in a balanced way without introducing massive amounts of misassemblies and errors.

Finally, detailed results of QUAST evaluation on the Human Chromosome 14 (HC14) dataset is summarized in Supplementary Table S3. Despite the highly repetitive nature of HC14 dataset as suggested in [8], three of the tools, Figbird, GapCloser and GapFiiler filled more than 30% nucleotides. From the overall comparison presented in Figure 5C, it can be observed that Figbird is second to GapCloser in terms of the amount of gap filled. But the additional 3% gap filled by GapCloser comes at the expense of 47% and 7% more misassemblies and erroneous length respectively compared to Figbird. Moreover, GapCloser fills smaller amount of gaps while making more misassemblies and errors than Figbird in the other datasets. On the other hand, Gap2Seq, which is able to reduce the gap length by the highest amount in the other two datasets, fills 16% less gaps than Figbird despite 30% more misassemblies.

Overall, our EM based method shows pareto optimal performance with respect to amount of gap filled, misassemblies and erroneous sequence introduced in all three datasets i.e. no other tool is able to fill more nucleotides while making less misassemblies or errors than Figbird. Figbird has achieved best or near best performance score for all the evaluation metrics in every dataset used in evaluation suggesting the usefulness of our approach of gap filling procedure.

### 3.4 Time and memory usage

We have compared the performance of Figbird in terms of run time and peak memory usage with four other tools in literature as mentioned in Section 3.1. The detailed commands and parameters used to run each of these tools can be found in Supplementary Note 3.3. Table3 shows the time and memory usage of the tools on the Human Chromosome 14 dataset. The comparisons on other two datasets are summarized on Supplementary Table S4 and S5. From Table 3, we can see that, GapCloser and Sealer are the two fastest tools in terms of run time across all the assemblies while Gap2Seq is the slowest among all. Figbird and GapFiller-bwa have moderate time requirements. In case of memory usage, Gap2seq and Sealer requires the highest amount of memory due to their problem formulation whereas GapFiller requires the lowest. The memeory requirement of Figbird is almost similar to that of GapCloser, taking less memory in case of Allpaths-LG, CABOG, MSR-CA etc. while taking slightly more memory in case of Velvet, ABySS etc. Overall, we find that although there are gap filling tools with lower time and memory usage than Figbird, it falls within the time and memory requirement range of the state-of-the-art tools. The increased run time can be regarded as a trade-off with the improved performance. Moreover, the time and memory requirements of Figbird scale linearly with the number and lengths of gaps, and the number of reads making it applicable to large datasets unlike some of the other gap filling tools.

**Table 3.**
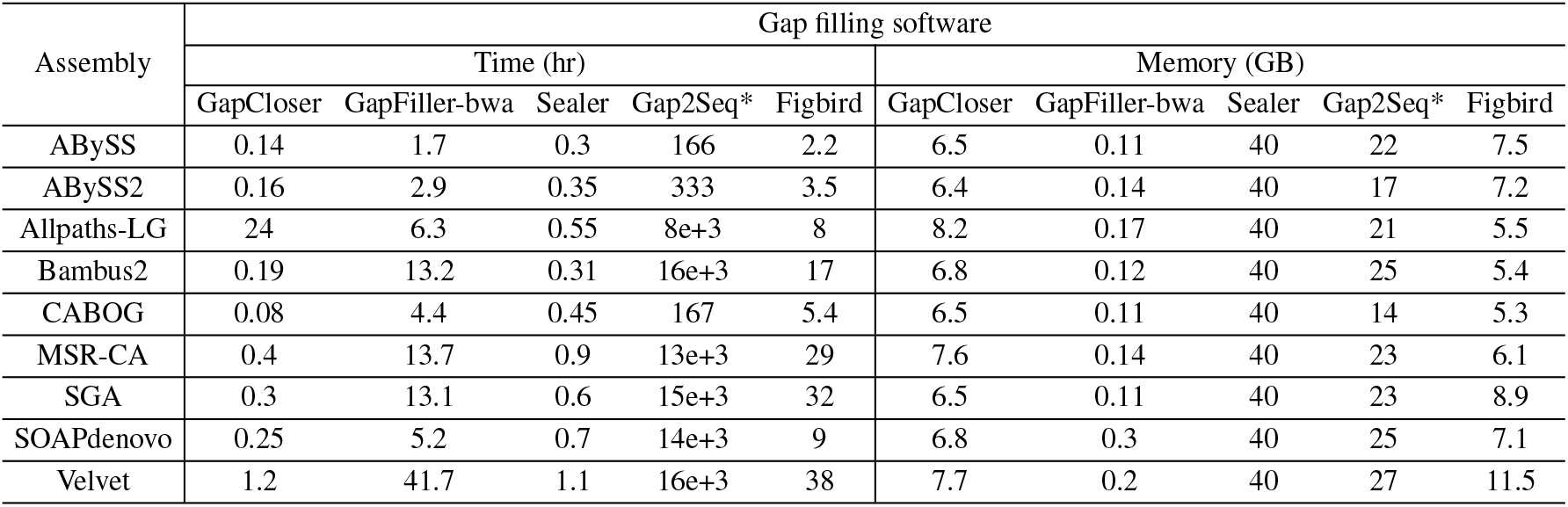
Gap-closing performance of all the tools on 9 draft genome assemblies of Human Chromosome14 genome.*The performance metric values for Gap2seq tool has been taken from the corresponding paper [10] as the tool could not be evaluated due to the high run time and memory constraints.

| Assembly | Gap filling software |  |  |  |  |  |  |  |  |  |
| --- | --- | --- | --- | --- | --- | --- | --- | --- | --- | --- |
|  | Time (hr) |  |  |  |  | Memory (GB) |  |  |  |  |
|  | GapCloser | GapFiller-bwa | Sealer | Gap2Seq* | Figbird | GapCloser | GapFiller-bwa | Sealer | Gap2Seq* | Figbird |
| ABYSS | 0.14 | 1.7 | 0.3 | 166 | 2.2 | 6.5 | 0.11 | 40 | 22 | 7.5 |
| ABYSS2 | 0.16 | 2.9 | 0.35 | 333 | 3.5 | 6.4 | 0.14 | 40 | 17 | 7.2 |
| Allpaths-LG | 24 | 6.3 | 0.55 | 8e+3 | 8 | 8.2 | 0.17 | 40 | 21 | 5.5 |
| Bambus2 | 0.19 | 13.2 | 0.31 | 16e+3 | 17 | 6.8 | 0.12 | 40 | 25 | 5.4 |
| CABOG | 0.08 | 4.4 | 0.45 | 167 | 5.4 | 6.5 | 0.11 | 40 | 14 | 5.3 |
| MSR-CA | 0.4 | 13.7 | 0.9 | 13e+3 | 29 | 7.6 | 0.14 | 40 | 23 | 6.1 |
| SGA | 0.3 | 13.1 | 0.6 | 15e+3 | 32 | 6.5 | 0.11 | 40 | 23 | 8.9 |
| SOAPdenovo | 0.25 | 5.2 | 0.7 | 14e+3 | 9 | 6.8 | 0.3 | 40 | 25 | 7.1 |
| Velvet | 1.2 | 41.7 | 1.1 | 16e+3 | 38 | 7.7 | 0.2 | 40 | 27 | 11.5 |

## 4 Conclusion

In the paper, we presented a probabilistic method based on the expectation-maximization (EM) algorithm to fill gaps in genome assemblies using paired-end and mate-pair reads from second generation sequencing technology with relatively low error-rate. As gap filling is one of the last stages in genome assembly pipeline and the comparatively easier regions of the genome have already been reconstructed by de novo assemblers in previous stages, only complex regions are left at this stage to fill up. So, the main objective was to incorporate essential information such as insert size distribution, sequencing errors, etc. to correctly estimate the true lengths of gaps and fill them with introduction of minimal amount of errors making downstream genome analysis easier. The results from experiments on multiple real datasets show that our method achieves a balanced performance across a variety of assembly pipelines managing to close a larger number of gaps, improving the overall NGA50 value while still maintaining to keep the amount of misassembly and length of erroneously introduced sequence low compared to existing. Specifically, it demonstrates pareto optimal performance in terms of the portion of gap filled and the numbers of misassemblies and errors introduced. An interesting future direction is to explore whether this approach can be extended to third generation sequencing data to accurately fill gaps that cannot be done using only second generation reads.

