## Supplementary Information for "Figbird: A probabilistic method for filling gaps in genome assemblies"

### 1 Supplementary Tables

#### 1.1 Detailed results obtained using QUAST

Table S1: Quality of the original and the gap-filled assemblies of the *Staphylococcus Aureus* genome using various tools.

| Tool | Original | Gap-Closer | GapFiller bowtie | GapFiller bwa | Gap2-Seq | Sealer | Figbird |
|---|---|---|---|---|---|---|---|
| **ABySS** | | | | | | | |
| Misassmblies | 5 | 40 | 0 | **-20** | 60 | 20 | 20 |
| Erroneous-length | 10587 | 26.3 | -3.5 | 31 | 70.5 | 32.5 | **-21.8** |
| Unaligned-length | 7935 | -21.9 | -10.2 | -10.2 | **-43** | 0 | -7.9 |
| NGA50 | 31079 | 0 | 0 | 0 | **0.3** | 0.3 | -1.2 |
| Number of gaps | 69 | -18.8 | -13 | -29 | **-87** | -71 | -29 |
| Total gap length | 55885 | -24.9 | -9.5 | -26 | **-94.5** | -59.4 | -17.6 |
| **ABySS2** | | | | | | | |
| Misassmblies | 5 | 20 | **0** | 40 | 40 | 40 | 20 |
| Erroneous-length | 10312 | -3.7 | 0.5 | 0.1 | -27.4 | -13.7 | **-30.9** |
| Unaligned-length | 0 | 0 | 0 | 0 | 0 | 0 | 0 |
| NGA50 | 106796 | 15.1 | 0 | 0 | **29** | 0 | 3.6 |
| Number of gaps | 35 | -34.3 | -11.4 | -31.4 | **-80** | **-80** | -57.1 |
| Total gap length | 9393 | -60.8 | -29.9 | -53.2 | **-94.5** | -93 | -79.9 |
| **Allpaths-LG** | | | | | | | |
| Misassmblies | 0 | **0** | 1 | 1 | **0** | **0** | **0** |
| Erroneous-length | 5991 | -22.7 | -5.9 | -8.4 | 9.7 | 55.5 | **-26.9** |
| Unaligned-length | 0 | 0 | 0 | 0 | 0 | 0 | 0 |
| NGA50 | 110168 | 2.7 | 35.9 | **69.6** | 48.5 | 35.9 | 31.5 |
| Number of gaps | 48 | -47.9 | 31.2 | -41.7 | **-70.8** | -68.8 | -52.1 |
| Total gap length | 9900 | -74.4 | -23.1 | -40.8 | **-94.7** | 86.6 | -72.9 |
| **Bambus2** | | | | | | | |
| Misassmblies | 0 | 1 | **0** | **0** | **0** | **0** | **0** |
| Erroneous-length | 24570 | **-30.6** | -4.4 | 16.9 | -1.4 | -19 | -15.6 |
| Unaligned-length | 0 | 0 | 0 | 0 | 0 | 0 | 0 |
| NGA50 | 40233 | **34.8** | 1.6 | 7.3 | 17.2 | 24 | 17.2 |
| Number of gaps | 99 | -67.7 | -14.1 | -18.2 | **-69.7** | -53.5 | -62.6 |
| Total gap length | 29205 | -78 | -23.9 | -37.2 | **-84.1** | 37.8 | -76.7 |
| **MSR-CA** | | | | | | | |
| Misassmblies | 10 | **-30** | **-30** | **-30** | -20 | 20 | **-30** |
| Erroneous-length | 17276 | -2.6 | 0.3 | 1.8 | -4 | **-12.3** | -9.1 |
| Unaligned-length | 0 | 0 | 0 | 0 | 0 | 0 | 0 |
| NGA50 | 64114 | 50.3 | 20.4 | 20.4 | 50.3 | 45.3 | **58.3** |
| Number of gaps | 81 | -51.9 | -19.8 | -29.6 | -56.8 | -50.6 | **-61.7** |
| Total gap length | 10353 | **-76.4** | -24.5 | -39.4 | -70.5 | -47.3 | -73.3 |
| **SGA** | | | | | | | |
| Misassmblies | 2 | 0 | 0 | 0 | **-50** | **-50** | **-50** |
| Erroneous-length | 13811 | -49.3 | -19.9 | -31.4 | -10.6 | -16.8 | **-53.8** |
| Unaligned-length | 0 | 0 | 0 | 0 | 0 | 0 | 0 |

*(continued)*

**Table S1 – Continued**

| Tool | Original | Gap-Closer | GapFiller bowtie | GapFiller bwa | Gap2-Seq | Sealer | Figbird |
|---|---|---|---|---|---|---|---|
| NGA50 | 9541 | 123.7 | 8.9 | 9.7 | 221.4 | 173.3 | **169.1** |
| Number of gaps | 654 | -75.1 | -20.8 | -37.3 | -80.1 | -78.7 | **-82.3** |
| Total gap length | 300607 | -54.3 | -5.7 | -10.1 | -72.1 | -61.6 | **-83.2** |
| **SOAPdenovo** | | | | | | | |
| Misassmblies | 2 | 0 | 0 | 0 | 0 | 0 | 0 |
| Erroneous-length | 35433 | -1.3 | 1.2 | .7 | -1.5 | -0.2 | **-2.4** |
| Unaligned-length | 4055 | **-100** | **-100** | **-100** | 3.9 | 0 | -7.5 |
| NGA50 | 69834 | 0 | 0 | 0 | 0 | 0 | 0 |
| Number of gaps | 9 | -22.2 | -22.2 | -33.3 | **-55.6** | -22.2 | -33.3 |
| Total gap length | 4857 | -60.4 | -24 | -30.8 | **-94.2** | -1.2 | -27.7 |
| **Velvet** | | | | | | | |
| Misassmblies | 25 | 8 | **0** | 4 | 8 | 12 | 4 |
| Erroneous-length | 24160 | -32.1 | -2.2 | -18.8 | **-36.3** | 13.8 | -15.6 |
| Unaligned-length | 1270 | **-49.4** | -20.5 | -21.3 | **-49.4** | 0 | **-49.4** |
| NGA50 | 46087 | 19.1 | 26 | 49 | **73.3** | 20.9 | 51.5 |
| Number of gaps | 128 | -46.9 | -30.5 | -41.4 | **-68.8** | -52.3 | -54.7 |
| Total gap length | 17688 | -59.6 | -37.9 | -49.2 | **-81.2** | -29.2 | -68.8 |
| **Total(average %)** | | | | | | | |
| Misassmblies | 49 | 5 | -3 | 0 | -3 | 6 | **-4** |
| Erroneous-length | 142140 | -14 | -4 | -1 | -2 | 5 | **-22** |
| Unaligned-length | 13260 | **-21** | -16 | -16 | -12 | 0 | -8 |
| NGA50 | 477852 | 31 | 12 | 20 | 51 | 37 | **65** |
| Number of gaps | 1123 | -45 | -20 | -32 | **-71** | -59 | -54 |
| Total gap length | 437888 | -61 | -22 | -36 | **-85** | -52 | -62 |

Table S2: Quality of the original and the gap-filled assemblies of the *Rhodobacter Sphaeroides* genome using various tools.

| Tool | Original | Gap-Closer | GapFiller bowtie | GapFiller bwa | Gap2-Seq | Sealer | Figbird |
|---|---|---|---|---|---|---|---|
| **ABySS** | | | | | | | |
| Misassmblies | 20 | **0** | **0** | **0** | 5 | 5 | **0** |
| Erroneous-length | 140634 | 1 | **-2.5** | 0.1 | -0.4 | -0.8 | -1.2 |
| Unaligned-length | 23522 | -7.2 | 100.9 | 70.8 | **-10** | **-10** | 106.6 |
| NGA50 | 6538 | 0.2 | 0.6 | 0.2 | **4.4** | 2.7 | 0.2 |
| Number of gaps | 323 | -20.1 | -8.7 | -19.8 | -45.8 | -39.6 | -36.5 |
| Total gap length | 114587 | -3.2 | -0.3 | -3.7 | **-19.6** | -8.6 | -5.6 |
| **ABySS2** | | | | | | | |
| Misassmblies | 12 | 8.3 | **0** | **0** | 133.3 | 0 | **0** |
| Erroneous-length | 15750 | 13.5 | **-0.4** | 0 | 38.5 | **-5** | 0.3 |
| Unaligned-length | 8230 | -0.4 | -1.9 | -36.1 | 0 | 0 | **-39.6** |
| NGA50 | 31197 | **11.1** | 0 | 3.8 | 11.1 | 5.7 | -12.1 |
| Number of gaps | 292 | **-21.2** | -1.4 | -5.8 | -19.9 | -10.3 | -11.3 |

*(continued)*

**Table S2 – Continued**

| Tool | Original | Gap-Closer | GapFiller bowtie | GapFiller bwa | Gap2-Seq | Sealer | Figbird |
|---|---|---|---|---|---|---|---|
| Total gap length | 62627 | -9.9 | 2.6 | -9 | **-38.7** | -8.7 | -6.3 |
| **Allpaths-LG** | | | | | | | |
| Misassmblies | 5 | 20 | **0** | **0** | **0** | **0** | **0** |
| Erroneous-length | 11738 | 97.1 | -2.6 | -2 | 3.4 | **-2.7** | 0.6 |
| Unaligned-length | 0 | 0 | 0 | 0 | 0 | 0 | 0 |
| NGA50 | 79634 | 11.5 | 2 | 0 | **12.8** | 0 | 0 |
| Number of gaps | 170 | -3.5 | -3.5 | -3.5 | -9.4 | -7.6 | **-10.6** |
| Total gap length | 21409 | -13.4 | 8 | 1.2 | **-25** | -10.2 | -10.7 |
| **Bambus2** | | | | | | | |
| Misassmblies | 5 | 0 | 0 | 0 | 0 | 0 | 0 |
| Erroneous-length | 106359 | -0.3 | -0.6 | -0.8 | 0.5 | **-4.2** | -0.5 |
| Unaligned-length | 4716 | -0.7 | -2.7 | -5.5 | **-100** | 0 | -11.5 |
| NGA50 | 15043 | 0 | 0 | 0 | **1.3** | 0.6 | 0.5 |
| Number of gaps | 85 | -15.3 | -5.9 | -7.1 | **-35.3** | -21.2 | -11.8 |
| Total gap length | 57041 | -14.2 | -9.6 | -13.6 | **-31.9** | -14.9 | -23.8 |
| **CABOG** | | | | | | | |
| Misassmblies | 15 | 0 | 0 | **-13.3** | **-13.3** | **-13.3** | 0 |
| Erroneous-length | 16750 | 44.1 | 0.3 | 0.4 | -1.9 | -1.4 | **-2.1** |
| Unaligned-length | 0 | 0 | 0 | 0 | 0 | 0 | 0 |
| NGA50 | 26819 | 0.8 | 11.4 | 11.4 | 3.9 | 3.9 | **24.8** |
| Number of gaps | 193 | -2.6 | -2.1 | -4.7 | -9.3 | -5.7 | **-18.1** |
| Total gap length | 21547 | -13 | 5.7 | -3.9 | **-22.5** | -7.3 | -10.5 |
| **MSR-CA** | | | | | | | |
| Misassmblies | **-20** | -20 | 0 | 10 | 270 | **-20** | -10 |
| Erroneous-length | 22522 | **-0.8** | 2.6 | 8.4 | 19.7 | **-0.9** | 1.5 |
| Unaligned-length | 1377 | 0 | 0 | 0 | 0 | 0 | 0 |
| NGA50 | 75776 | -5.3 | 19 | 20 | 13.9 | -1.1 | **44** |
| Number of gaps | 356 | -12.1 | -7.3 | -10.4 | -26.4 | -5.3 | **-29.5** |
| Total gap length | 32628 | -19.9 | 3.4 | -7 | **-67.1** | -9.8 | -27.3 |
| **SGA** | | | | | | | |
| Misassmblies | 2 | 0 | 0 | 0 | 1600 | 0 | 50 |
| Erroneous-length | 58135 | 4.1 | -2.9 | -5.1 | 33.9 | **-13.3** | -1.3 |
| Unaligned-length | 69226 | -0.7 | -12.5 | -13.2 | **-41** | -24.4 | -24.3 |
| NGA50 | 2601 | 5.7 | 1.2 | 5.2 | **98** | 31.8 | 8.6 |
| Number of gaps | 938 | -8.6 | -3.9 | -7.7 | **-37.1** | -18.8 | -13.2 |
| Total gap length | 1145600 | -2.2 | -0.3 | -2.7 | **-23.3** | -9.4 | -10.4 |
| **SOAPdenovo** | | | | | | | |
| Misassmblies | 3 | **0** | **0** | **0** | 33.3 | 0 | **0** |
| Erroneous-length | 56228 | 8.7 | -0.1 | 0.1 | **-7.1** | -0.5 | 0.9 |
| Unaligned-length | 0 | 0 | 0 | 0 | 0 | 0 | 0 |
| NGA50 | 27434 | **0** | -1.2 | -1.2 | **0** | 0 | -1.2 |
| Number of gaps | 38 | 0 | 0 | 0 | **-10.5** | -2.6 | **-10.5** |
| Total gap length | 10461 | -10 | 2.4 | 0.3 | **-20.2** | -3.5 | -17.3 |
| **Velvet** | | | | | | | |
| Misassmblies | 19 | 15.8 | 0 | -15.8 | 10.5 | 0 | **-26.3** |

*(continued)*

**Table S2 – Continued**

| Tool | Original | Gap-Closer | GapFiller bowtie | GapFiller bwa | Gap2-Seq | Sealer | Figbird |
|---|---|---|---|---|---|---|---|
| Erroneous-length | 40419 | **-14.5** | -4.5 | 3.2 | -5.4 | -5.5 | 10.9 |
| Unaligned-length | 28344 | -3 | -5.9 | -7.6 | -17.4 | -1.5 | **-22.7** |
| NGA50 | 54238 | **0.3** | -9.8 | -0.9 | 0 | 0 | -16.4 |
| Number of gaps | 427 | -11.2 | -5.4 | -9.4 | -21.5 | -13.3 | **-29.3** |
| Total gap length | 86815 | -7.6 | 0 | -6.3 | **-26.3** | -3.8 | -13.3 |
| **Total(average %)** | | | | | | | |
| Misassmblies | 91 | 3 | 0 | **-2** | 227 | -3 | 2 |
| Erroneous-length | 468535 | 18 | **-1** | 1 | 10 | -3 | 2 |
| Unaligned-length | 135415 | -1 | 9 | 1 | **-18** | -3 | -5 |
| NGA50 | 319280 | 2 | 3 | 5 | **17** | 5 | 6 |
| Number of gaps | 2822 | -10 | -4 | -7 | **-23** | -13 | -18 |
| Total gap length | 1552715 | -10 | 2 | -4 | **-30** | -8 | -13 |

Table S3: Quality of the original and the gap-filled assemblies of Human Chromosome 14 using various tools.

| Tool | Original | Gap-Closer | GapFiller bowtie | GapFiller bwa | Gap2-Seq | Sealer | Figbird |
|---|---|---|---|---|---|---|---|
| **ABySS** | | | | | | | |
| Misassmblies | 3 | 133 | 33.3 | **0** | 100 | 66.7 | **0** |
| Erroneous-length | 190458 | 18.2 | 7.6 | 7.4 | -9.4 | **-17.6** | 8.5 |
| Unaligned-length | 262068 | -16.6 | -28.9 | **-34.1** | -8.4 | -3.6 | -19.5 |
| NGA50 | 1320 | 1 | 0.5 | 0.7 | **1.3** | 1.2 | 0.8 |
| Number of gaps | 1061 | -5.9 | -28.9 | -32.5 | **-33.4** | -46.6 | -27.2 |
| Total gap length | 585628 | -24.5 | -23.4 | **-27.5** | -25.5 | -25.6 | -22.9 |
| **ABySS2** | | | | | | | |
| Misassmblies | 99 | 18.2 | **2** | 4 | 5.1 | 0 | 8.1 |
| Erroneous-length | 555099 | 15.9 | 1.5 | 2.8 | 3.5 | **-6.5** | 3.1 |
| Unaligned-length | 157759 | -21.4 | -28.8 | **-36.3** | -15.4 | -2.2 | -26.4 |
| NGA50 | 11869 | **4.1** | 1.5 | 3 | 2.4 | 2.9 | 3.2 |
| Number of gaps | 2820 | -14.5 | -29.5 | -38.5 | -23.9 | -22.69 | **-43.8** |
| Total gap length | 949137 | -34.9 | -10.6 | -25.3 | **-36.8** | -15.5 | -28.1 |
| **Allpaths-LG** | | | | | | | |
| Misassmblies | 95 | **-6.3** | 5.3 | 8.4 | 14.7 | 2.1 | 6.3 |
| Erroneous-length | 667229 | 34.5 | -0.6 | 7.8 | **-3** | -3.4 | 8.6 |
| Unaligned-length | 36941 | **-14.3** | -11.1 | -11.9 | 26.8 | 12.3 | 3.8 |
| NGA50 | 34534 | **48.3** | 20.7 | 23 | 23.3 | 6 | 32.5 |
| Number of gaps | 4307 | **-35.1** | -19.3 | -20.6 | -29.8 | -17.7 | -33.6 |
| Total gap length | 3227193 | -37.9 | -16 | -17.2 | -16 | -7.2 | **-39.4** |
| **Bambus2** | | | | | | | |
| Misassmblies | 1584 | 3.1 | 2.3 | 5.4 | **2.1** | -0.2 | 3.4 |
| Erroneous-length | 11114542 | **-9.6** | -0.2 | 14.8 | -0.6 | -18.2 | -3.9 |
| Unaligned-length | 161358 | **-42.7** | -35.8 | -31.3 | 2.5 | -33.8 | -37.3 |

*(continued)*

**Table S3 – Continued**

| Tool | Original | Gap-Closer | GapFiller bowtie | GapFiller bwa | Gap2-Seq | Sealer | Figbird |
|---|---|---|---|---|---|---|---|
| NGA50 | 3045 | **34.8** | 12 | 15.2 | 1.8 | 23.7 | 25.1 |
| Number of gaps | 11809 | **-16.4** | -2.3 | -2.3 | -6.6 | -22.3 | -1.6 |
| Total gap length | 10370362 | -45.6 | -18.9 | -29.3 | -4.6 | -21.4 | **-46.6** |
| **CABOG** | | | | | | | |
| Misassmblies | 91 | 16.5 | 7.7 | 5.5 | 7.7 | **2.2** | 4.4 |
| Erroneous-length | 615239 | 19 | -2.2 | 0.2 | -3.5 | -4.4 | -0.3 |
| Unaligned-length | 2506 | 0 | 0 | 0 | 0 | 0 | 0 |
| NGA50 | 46665 | 16.2 | 58.8 | **62.3** | 8.9 | 11.9 | 39.8 |
| Number of gaps | 3043 | -18.9 | -46.5 | **-51.2** | -13.8 | -15.1 | -38.6 |
| Total gap length | 231078 | **-50.3** | -34.9 | -41.2 | -22.6 | -19.9 | -32.5 |
| **MSR-CA** | | | | | | | |
| Misassmblies | 1110 | 14.5 | 9.9 | 25.3 | 6.8 | **1.3** | 13.3 |
| Erroneous-length | 5412965 | 2.8 | 3.9 | 21.7 | **-6** | -6.5 | -4.9 |
| Unaligned-length | 318421 | -30.5 | -28.8 | -33 | -13.1 | -9 | **-31.2** |
| NGA50 | 5704 | 73.3 | 65.9 | **77.7** | 32.9 | 18.9 | 52.5 |
| Number of gaps | 30622 | -34.9 | -42.4 | **-49.5** | -27.8 | -26 | -46.5 |
| Total gap length | 6097928 | **-49.3** | -37.8 | -49.2 | -18.1 | -11.3 | -48.3 |
| **SGA** | | | | | | | |
| Misassmblies | 8 | 375 | 37.5 | 87.5 | 287.5 | **12.5** | 150 |
| Erroneous-length | 1580489 | 21.1 | -18.4 | -13.9 | -24.6 | **-26.6** | -0.6 |
| Unaligned-length | 1160159 | -83.9 | -82.8 | -87 | -38.6 | -22.1 | **-90.5** |
| NGA50 | 2644 | **244.2** | 206.6 | 239.3 | 149.1 | 80.4 | 164.6 |
| Number of gaps | 21459 | **-56.7** | -46.3 | -49.9 | -51.5 | -39.4 | -49 |
| Total gap length | 12840408 | -53.5 | -50 | -55.4 | -30.2 | -17.6 | **-62.9** |
| **SOAPdenovo** | | | | | | | |
| Misassmblies | 1250 | 17.1 | 3.5 | 9.4 | 11.7 | **1.6** | 13.1 |
| Erroneous-length | 8449941 | -1.3 | 1.1 | 3.2 | -0.5 | **-4.7** | 0.2 |
| Unaligned-length | 1306173 | **-28.8** | -24.4 | -28.1 | -14.1 | -10.1 | -27.2 |
| NGA50 | 6592 | **17.4** | 3.6 | 4.1 | 4 | 7.5 | 4.9 |
| Number of gaps | 8544 | **-25.2** | -4.5 | -5.1 | -18.8 | -11.7 | -5.7 |
| Total gap length | 10255930 | **-21.3** | -11.3 | -15.9 | -6.3 | 6 | -18 |
| **Velvet** | | | | | | | |
| Misassmblies | 9308 | 26.3 | 24.8 | 51.5 | 13.3 | **10.5** | 24 |
| Erroneous-length | 12531431 | -10.4 | 32.3 | 58.6 | -0.5 | **-23.1** | 23.5 |
| Unaligned-length | 23484076 | -58.4 | -54.9 | **-70** | -20.7 | -31 | -52.4 |
| NGA50 | 1793 | **104.4** | 49.6 | 67.5 | 27 | 65.1 | 43.1 |
| Number of gaps | 51567 | **-43.4** | -24.1 | -26.6 | -26.6 | -37 | -21.5 |
| Total gap length | 63559964 | **-22.8** | -14.7 | -22.8 | -4.2 | -11.8 | -17.1 |
| **Total(average %)** | | | | | | | |
| Misassmblies | 13449 | 67 | 15 | 22 | 50 | **11** | 20 |
| Erroneous-length | 40562294 | 11 | 3 | 12 | -4 | **-12** | 4 |
| Unaligned-length | 26731702 | -32 | -32 | **-36** | -9 | -11 | -34 |
| NGA50 | 102297 | **61** | 47 | 55 | 28 | 25 | 40 |
| Number of gaps | 132412 | -27 | -27 | **-30** | -25 | -26 | -29 |
| Total gap length | 107168491 | **-37** | -24 | -31 | -18 | -15 | -34 |

#### 1.2 Time and memory usage

Table S4: Gap-closing performance of the tools mentioned in the paper on 8 draft genome assemblies of *Staphylococcus aureus*.

| Assembly | Software | Time(min) | Memory(GB) |
|---|---|---|---|
| ABySS | GapCloser | 0.2 | 0.68 |
| | GapFiller-bwa | 5.8 | 0.12 |
| | Sealer | 7.1 | 40 |
| | Gap2seq | 0.2 | 1.95 |
| | Figbird | 10 | 0.7 |
| ABySS2 | GapCloser | 0.23 | 0.67 |
| | GapFiller-bwa | 5.6 | 0.15 |
| | Sealer | 7.5 | 40 |
| | Gap2seq | 0.2 | 1.95 |
| | Figbird | 16 | 0.72 |
| Allpaths-LG | GapCloser | 2.2 | 0.71 |
| | GapFiller-bwa | 5.6 | 0.12 |
| | Sealer | 7.2 | 40 |
| | Gap2seq | 0.3 | 1.95 |
| | Figbird | 39 | 0.71 |
| Bambus2 | GapCloser | 0.23 | 0.72 |
| | GapFiller-bwa | 7.5 | 0.12 |
| | Sealer | 8.1 | 40 |
| | Gap2seq | 0.45 | 1.9 |
| | Figbird | 63 | 0.75 |
| MSR-CA | GapCloser | 0.26 | 0.73 |
| | GapFiller-bwa | 6.6 | 0.11 |
| | Sealer | 8.5 | 40 |
| | Gap2seq | 0.65 | 1.92 |
| | Figbird | 34 | 0.79 |
| SGA | GapCloser | 0.15 | 0.73 |
| | GapFiller-bwa | 15.2 | 0.15 |
| | Sealer | 8.6 | 40 |
| | Gap2seq | 2 | 1.9 |
| | Figbird | 77 | 0.9 |
| SOAPdenovo | GapCloser | 0.16 | 0.69 |
| | GapFiller-bwa | 4.7 | 0.11 |
| | Sealer | 6.5 | 40 |
| | Gap2seq | 0.3 | 1.9 |
| | Figbird | 10 | 0.7 |
| Velvet | GapCloser | 0.17 | 0.71 |
| | GapFiller-bwa | 7.5 | 0.12 |
| | Sealer | 8.1 | 40 |
| | Gap2seq | 0.9 | 1.9 |
| | Figbird | 35 | 0.8 |

Table S5: Gap-closing performance of the tools mentioned in the paper on 9 draft genome assemblies of *Rhodobacter sphaeroides*.
*The performance metric values for Gap2seq tool has been taken from their paper [1] as the tool could not be evaluated due to the high run time and memory constraints.

| Assembly | Software | Time(min) | Memory(GB) |
|---|---|---|---|
| ABySS | GapCloser | 0.1 | 0.41 |
| | GapFiller-bwa | 17 | 0.11 |
| | Sealer | 7.2 | 40 |
| | Gap2seq* | 800 | 2.1 |
| | Figbird | 22 | 0.78 |
| ABySS2 | GapCloser | 0.11 | 0.42 |
| | GapFiller-bwa | 21.3 | 0.12 |
| | Sealer | 7.1 | 40 |
| | Gap2seq* | 998 | 2.1 |
| | Figbird | 25 | 0.8 |
| Allpaths-LG | GapCloser | 10.9 | 0.82 |
| | GapFiller-bwa | 7.3 | 0.12 |
| | Sealer | 7.3 | 40 |
| | Gap2seq* | 578 | 2.2 |
| | Figbird | 19 | 0.79 |
| Bambus2 | GapCloser | 0.35 | 0.61 |
| | GapFiller-bwa | 10.1 | 0.14 |
| | Sealer | 7.5 | 40 |
| | Gap2seq* | 991 | 2.1 |
| | Figbird | 176 | 0.83 |
| CABOG | GapCloser | 0.2 | 0.55 |
| | GapFiller-bwa | 13.8 | 0.11 |
| | Sealer | 7.2 | 40 |
| | Gap2seq* | 121 | 2.2 |
| | Figbird | 23 | 0.8 |
| MSR-CA | GapCloser | 0.25 | 0.64 |
| | GapFiller-bwa | 20.8 | 0.11 |
| | Sealer | 7.8 | 40 |
| | Gap2seq* | 858 | 2.5 |
| | Figbird | 68 | 0.8 |
| SGA | GapCloser | 0.2 | 0.67 |
| | GapFiller-bwa | 30.1 | 0.15 |
| | Sealer | 7.8 | 40 |
| | Gap2seq* | 10e+3 | 2.1 |
| | Figbird | 91 | 1.1 |
| SOAPdenovo | GapCloser | 0.3 | 0.59 |
| | GapFiller-bwa | 2.4 | 0.12 |
| | Sealer | 7.8 | 40 |
| | Gap2seq* | 105 | 2.1 |
| | Figbird | 10 | 0.8 |

| | | | |
|---|---|---|---|
| Velvet | GapCloser | 0.1 | 0.58 |
| | GapFiller-bwa | 13.1 | 0.12 |
| | Sealer | 7.1 | 40 |
| | Gap2seq* | 1221 | 2.1 |
| | Figbird | 54 | 1.1 |

### 2 Supplementary Figures

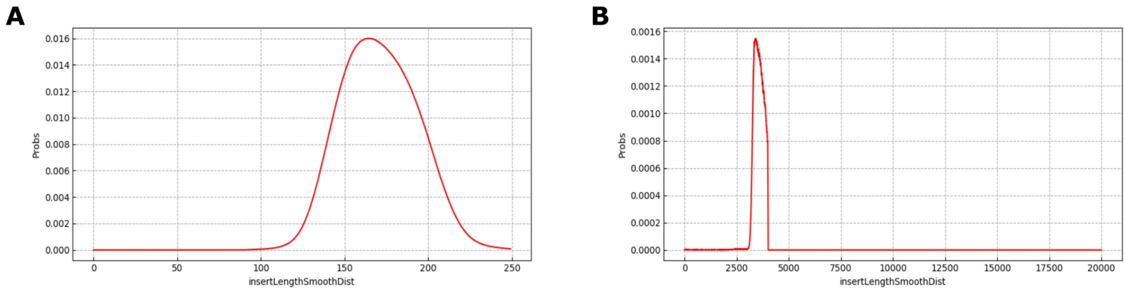

Figure S1: Insert size probabilities learned for (A) frag library and (B) jump library on ABySS2 assembly of *Staphylococcus Aureus* bacterial genome.

### 3 Supplementary Notes

#### 3.1 Script iterations

To run our method as a software Figbird, we have prepared a script to manage the entire program. Some parameters, details of iterations and types of reads, and some checking performed during the process are discussed below.

The read pairs are used in following modes for gap filling:

| Read type | Read library |
|---|---|
| Partial | Fragment |
| Unmapped | Both fragment and jump |

Table S6: Reads used in gap filling

As, fragment reads had higher length compared to jump reads, we used only this read as partial reads as there is a higher chance of partially falling in gaps for these reads. Also as longer reads contain more N, only the reads with N count less than or equal to 3 is considered. All the reads are checked so that they don't contain any character other than A,C,G,T and N. We have also considered gaps with length 1 unlike other methods. Sometimes filling a 1 length gap can increase or reduce erroneous length and NGA50 by a huge margin because of incorrect gap length estimation during the scaffold assembly. That's why although these gaps were ignored in other methods, we have tried to fill every possible gap.

Our method is implemented in an iterative fashion where multiple iterations are performed using different sets of reads with different types. The reason for an iterative approach is that a lot of gaps that have been filled in previous iterations can act as mapping regions for the reads that are currently unmapped. So a lot of reads that were unmapped in first few iterations will now have a higher chance of getting mapped and thus we can extract more out of the higher insert size jump libraries. For each iteration, the alignment using Bowtie2 is done twice. This is to find the partially aligned reads initially in the local alignment mode and then to find the one end unmapped reads. As fragment type library has much higher number of read pairs (36 million read pairs as opposed to 14 million in case of jump library for *Staphylococcus aureus* genome, we reduced fragment read set after first iteration by removing those pairs whose both end has been mapped (Cigar string of 101M in SAM file), so that subsequent iterations have less read to map.

In iteration 1, we use fragment reads as partial reads (soft clipped) and initialize probabilities with soft clipped fragment type reads and use these partial reads to fill the gaps. In iteration $2-3$, we use jump reads as unmapped reads and initialize probabilities with soft clipped frag type reads. The reason for using jump reads at the beginning part of the iterations is because of their comparative higher insert size. So, they were able to close more gaps and larger gaps at the beginning, that means the burden will be less for later iterations with less and small length gaps. In iteration 4 and 5, we use fragment reads as partial reads and jump type reads as unmapped reads to fill the gaps respectively. Then finally, in iteration $6-8$, we use frag reads as soft clipped reads and initialize probabilities with soft clipped frag type reads to complete the gap filling process. The reason for using these three iterations at ending part is that there are a lot of gaps which have small flanking regions on either side. So, using a library with higher insert are not useful in this scenario and thus the choice of frag type reads is made which has lower insert size.

At any point during the run of Figbird, if there is no change in gap length for the current iteration compared to previous one, we will break the loop and end the process there. To make these iterations faster, we will reduce the read set as well as the scaffold set i.e. only those scaffolds having gaps is kept and rest are removed. While placing the reads in gaps, we have chosen insert size threshold to be 3 times the left and right standard deviations from the mean calculated from the fully mapped read pairs for that library of reads. Finally the number of EM iterations during gap filling were kept 3 for partially aligned reads and 200 for the case of unmapped reads, with a break condition from the loop if the consensus is complete before the limit or it is stuck for 5 iterations straight.

#### 3.2 Adjustment of gap filling based on heuristics

**Exploration of range for gap estimates likelihood calculation:**
As discussed in the methodology section of paper, for each gap with length $l_g$, we will compute $l(\mathcal{G};\mathcal{R})$ for all $\mathcal{G}$ with length between $0.5 \times l_g$ to $2.5 \times l_g$. This will be done for gaps with length $\leq 400$. Otherwise we will fill the gaps considering their original length. But gap lengths given are not always exactly accurate. So to avoid the problem of filling gaps exactly with the same length, we do the following:

```
NNNNNNNNNNNNNNNNNNNNNNNNNNNNNNNNNNNNNNNNN
ACGTACGTNNNNNNNNNNNNNNNNNNNNNNNNNNNACGTACGT
ACGTACGTACGTACGT......................ACGTACGTACGTACGTACGTT
```

Figure S2: Consensus string at each EM iteration

Gaps are iteratively filled at each iteration as shown in Figure S2. If the number of N in consensus sequence becomes less than two times the read length, then we stop EM iteration and break from the loop. As the gap length became smaller, it will be evaluated as the range specified above in next iterations. Also, in the finalize step, for this type of gaps, we further cut read length characters from either side to make sure it gets evaluated using the range.

**Detection of repetitive region:** Sometimes, in our gap filling process, we will skip some gaps from filling process based on some repeat detection condition. This is only applicable in case of read pairs with low insert size($< 250$ bp) i.e. fragment type reads (partial or unmapped) in our case. To detect such gap, what we do is following: For each read, we evaluate the following 2 conditions:

- If a chunk of characters ($\geq 20$) characters from left side of the gap start position is present in the read more than 1 time
- If a chunk of characters ($\geq 20$) characters from right side of the gap start position is present in the read more than 1 time

If both of this conditions are met for the same read, then the region is considered as highly repetitive and we don't fill that gap. Otherwise, if one of the above conditions is met and and original gap is greater than 6 times (a high value is given to make sure the gap length surpasses the insert size) the read length, the gap is not filled. The reason for doing so is that as it is a large gap and insert size for this condition is lower, then filling it with partial or unmapped frag does not help. As we have to depend on jump library for these cases, so we avoid them from filling up incorrectly as jump reads perform better among the two libraries in these cases.

**Negative overlap detection:** Sometimes there are some gaps present in scaffold where, there shouldn't have been a gap. That means the gap should be closed down and there is an indirect overlap between the left and right flank sequences of these gaps, which we are calling the negative overlap. We have considered this for gaps with length $\leq 30$ (default value) which can be set by user. If the two flank sequences have an overlap greater than 5 bp and if there is any read with sufficiently long length that supports this merging of sequences, then we remove the gap and merge both sides into one and shrink the gap. The number of such gaps is very low (5 out of 1000) and only present in some assembly such as MSR-CA and SGA.

**Maximum likelihood modification:** Our maximum likelihood calculation method is modified a little based on type of reads we are using i.e. partial or unmapped to account for various erroneous scenarios during read placement. They are discussed below:

**Likelihood modification for invalid reads:** While placing a read in the gap, if it falls within the insert size range allowed, we place the read in certain range in all possible positions and calculate the best position of that read using the likelihood computed using the model parameters and probability distributions. Finally we add this probability to calculate the sum of likelihood for that gap estimate. Now, to solve the unnecessary read problem and incorrect gap length prediction specially in case of jump reads, we do the following: We compute the consensus sequence based on above placement of reads and for each read, find the probability based on sequence similarity and 3 other distributions- error, insertion, deletion distributions by read position and compare it with gap probability cutoff value. If probability is less than log(cutoff value), we accept this likelihood value in sum calculation, otherwise we add a penalty for that read (-50). The reason to put a fixed negative penalty is to discard a gap estimate of smaller length and stop shrinking the gap. If there are more valid reads falling in the gap, there will be more used reads, thus gap estimate will be close to actual one.

**Likelihood modification based on coverage:** We also modify our sum likelihood calculation based on gap coverage as shown in figure below:

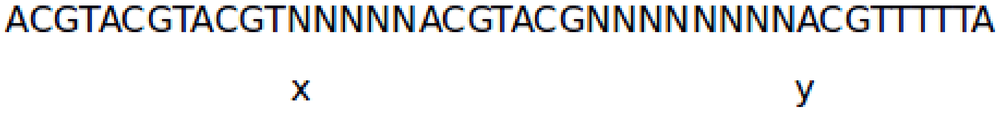

Figure S3: Consensus string at each EM iteration

In case of such scenarios, we try to find the starting and ending position of such fragmented regions i.e. x and y in above figure and discard all the reads that fall in this region by adding a similar penalty (-50) sum of max likelihood calculation. Note that they are mostly the unnecessary reads as the correct reads tend to have overlap.

**Likelihood modification based on the overlapping characteristics:** Based on the overlapping characteristics of all the reads either unmapped or partially assembled, a penalty scheme is added. The penalty scheme for unmapped reads contains the followings:

- Gap penalty: Gap found between reads aligned to the left flank and right flank and going inwards.
- Overlap threshold penalty: If overlap between the reads is less than 4 bp for that particular read placement.
- Left/ right alignment advantage: A likelihood boost based on the count of the reads that has more than 4 character aligned to the left or right side of the gap.

In case of partially mapped reads, our overlapping characteristics are determined as follows. For each gap estimate, we call a function that detects whether there is a overlap between reads and does the following. If there is a read that covers the entire gap and matches with both left and right regions of the gap sequence with fixed low threshold of error, we add a positive value to the maximum likelihood. Otherwise, if there is a correct overlap with reads from both sides,then we find the overlap length and add positive value to maximum likelihood based on that count of overlaps. Else if there is a false overlap detected, then we give a penalty for that gap estimate.

**Consensus probability update based on intermediate alignment:** In an ideal scenario, gaps are filled from one/both flanks and eventually merges at some EM iteration. But sometimes the consensus sequence gets updated circularly i.e. it changes back and forth into the same sequence every other iteration or it gets stuck as the EM iteration carries on at one/both sides as shown in figure below:

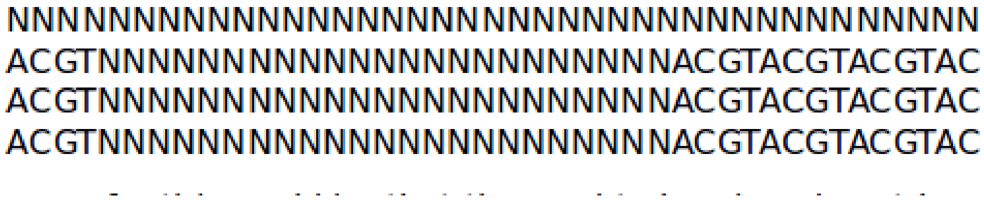

Figure S4: Consensus string gets stuck after certain EM iterations

The reason for this is that the read to be placed next has very little overlap with the filled consensus sequence and thus our algorithm can't find the correct position to place it. The advantage of this process is not only to solve these issues but also to make the convergence process faster. EM is by default a slow algorithm and the probabilities take a long time to converge. So, to solve these problems, we will perform the update if a combination of conditions are met. They are:

- If the consensus is stuck for more than two iterations.
- If the length of the gap is $\geq 400$ and thus for this gap estimate we didn't run a range of gap estimate calculation.

If both of these conditions are met, we do the following for each flank. We find those reads that has a certain threshold of base pair match with current filled consensus sequence of that flank. Conditions for accepting such read are that they are not yet accepted or placed, the read is in proper insert size range, segment taken from left or right flank must be greater than 20 and match count between prefix and suffix depends on the length of the read. Then finally, we update the probability with the consensus of all such reads considered.

**Clearing a filled sequence based on threshold values:** Sometimes, because of wrong read placement due to erroneous reads or false alignment by Bowtie2, some gaps can be wrongly filled by our method. We have tried to find such filled gaps by performing a number of checks to make sure that the gap sequence predicted from the consensus is consistent with the gap length and left or right flank based on multiple conditions specified below otherwise clear them to their original length to avoid misassemblies. For all these checks, we have considered an overlap threshold of 4. The conditions of clearing such gaps are:

- If the amount of left flank aligned character or right flank aligned character in consensus is less than overlap threshold then clear that particular sided aligned reads and build a new consensus.
- If there is no aligned read with left and/or right flank such as shown below, then clear the gap.

```
NNNNNNNNNNNNNNNNNNNNNNNNNNNNNNNNNNNNNNNNNNNNNNNNNNNNNNNN
  ACGATACGATACGATACGAT            ACGATACGATACGATACGAT
       ACGATACGATACGATACGAT            ACGATACGATACGATACGAT
```

Figure S5: Zero alignment with both flanks

- There will be a check for discontinuity among the placed reads. In case there is a pair of read found, where the overlap between the two reads is less than or equal to 2, it will be considered as a fragmented or discontinuous sequence and a read length amount character is going to be chopped off from the point of discontinuity to outwards direction in both sides.

```
      NNNNNNNNNNNNNNNNNNNNNNNNNNNNNNNNNNNNNNNNNNNNNNNNNNNNNNNN
ACGATACGATACGATACGAC
                  ACGATACGATACGATACGAT
                                    ATGATACGATACGATACGAT
```

Figure S6: An illustration of discontinuous placement of reads

#### 3.3 Command and parameter used to run the tools

To evaluate using GapCloser and GapFiller, we have provided a separate configuration file for each of them during each assembly run. Following are the commands with specific parameters used to run for each tool. All other parameters that are not mentioned in commands have been kept default.

- GapCloser:
  ./GapCloser -t 12 -l 101 -a scaffold.fa -b dataset.config -o output.fa
- GapFiller:
  perl GapFiller.pl -l dataset.config -s scaffold.fa -i 10 -T 24 -b output.fa
- Gap2Seq:
  ./Gap2Seq –scaffolds genome.scf.fasta –filled genome.scf.fill.fasta –reads frag1.fastq frag2.fastq,shortjump1.fastq,shortjump2.fastq

- Sealer:
./abyss-sealer -b40G -k90 -k80 -k70 -k60 -k50 -k40 -k30 -o output -S scaffold.fa -j 24 -P 10 -B 3000 -F 4000 frag1.fasta frag2.fasta shortjump1.fasta shortjump2.fasta

- Figbird:
./RunFigbird.sh Config.json > output.txt

  In the configuration files for GapCloser, GapFiller and Figbird, we have given average insert size as 180 for fragment library in case of *S.aureus* and *R. sphaeroides* dataset as input. In case of Human Chromosome14 we have used 160. For jump library, the insert size is 3500 for *S.aureus* and *R. sphaeroides* dataset and 2500 for Human Chromosome14 dataset for all three above mentioned tools The standard deviation in GapFiller tool has been given 0.25 for all three datasets.
